# Multidimensional phenotyping predicts lifespan and quantifies health in *C. elegans*

**DOI:** 10.1101/681197

**Authors:** Céline N. Martineau, André E. X. Brown, Patrick Laurent

## Abstract

Ageing affects a wide range of phenotypes at all scales, but an objective measure of ageing remains challenging, even in simple model organisms. We assumed that a wide range of phenotypes at the organismal scale rather than a limited number of biomarkers of ageing would best describe the ageing process. Hundreds of morphological, postural and behavioural features are extracted at once from high resolutions videos. A quantitative model using this multi-parametric dataset can predict the biological age and lifespan of individual *C. elegans*. We show that the quality of predictions on a held-out data set increases with the number of features added to the model, supporting our initial hypothesis. Despite the large diversity of ageing mechanisms, including stochastic insults, our results highlight a robust ageing trajectory, but variable ageing rates along that trajectory. We show that healthspan, which we defined as the range of abilities of the animals, is correlated to lifespan in wild-type worms.

## Introduction

Ageing is a plastic process altering phenotypes at the molecular, cellular, tissue and organism levels. Ultimately these alterations affect longevity and health of the organism. *C. elegans* has been proven a powerful model organism to identify molecular pathways regulating longevity (Kenyon, 2010). Several studies also used this organism to assess healthspan (as the period of life when animals are in good health) and compare short-lived and long-lived animals after video tracking (Bansal et al., 2015; Hahm et al., 2015; Zhang et al., 2016). Ideally suited for longitudinal studies, the morphological and behavioural repertoire of the worm offers numerous easily quantifiable parameters that can be measured non-invasively. Importantly, the behaviour of *C. elegans* evolves from the first day of adulthood as a consequence of modified neuronal functions and as a consequence of the stochastic senescence of muscle cells (Herndon et al., 2002; Liu et al., 2013). Hahm *et al*. used a single feature, maximum velocity, as an indicator of health, and showed that a *daf-2* mutation improves healthspan, and therefore couples lifespan and healthspan extensions (Hahm et al., 2015), in contrast to the previous results obtained by Bansal *et al*. (Bansal et al., 2015). On the other hand, Zhang *et al*. used a set of 5 biomarkers of ageing to conclude that the most plastic period of ageing is the end of life and that long-lived individuals have a longer span of poor health (Zhang et al., 2016).

Similarly to Zhang *et al*. who used several biomarkers of ageing to predict prognosis (Zhang et al., 2016), we postulated that a large set of phenotypic features would better cover the wide phenotypic evolution occurring during ageing, and therefore allow us to define better predictors or combinations of predictors of age and lifespan. Hundreds of morphological and behavioural features extracted at once from high resolution videos of worms was previously shown to produce meaningful classes of mutants (Javer et al., 2018; Javer Avelino et al., 2018; Yemini et al., 2013). We opted for this approach to build a multiple parameter database describing the phenotypic progression of worms during ageing.

Following previous work (Pincus and Slack, 2010), we define a biomarker of ageing as any phenotype that correlates with relative age. By this definition, we identified a set of 837 biomarkers of ageing. We used appropriate combinations of phenotypic features to predict biological age, prognosis and lifespan. We show that short-lived and long-lived animals follow the same trajectory of ageing and spend the same relative time in good health, according to our health index.

## Results

### Extreme phenotypes correlate with age and relative age

To assess the physiology of freely moving animals throughout their entire lives, hundreds phenotypes were measured from video tracking of individual worms. In total, the phenotypes of 151 wild-type worms (N2, Bristol) were followed every day of their lives from the L4 stage. From these videos, 1019 features are extracted representing morphological, postural and locomotion features displayed by the worms during recording. Similarly to the short physical performance battery (SPPB) test used in humans (Guralnik et al., 1994), and the maximum velocity in *C. elegans* (Hahm et al., 2015), the best markers of health might be extremes values of physical performance. Therefore the phenotypes were measured in basal conditions and after mechanical stimulation to extract basal and stimulated values for each of the measured features. For each of the 1019 tested features, Pearson’s and Spearman’s correlation coefficients with age and relative age are globally higher after the mechanical stimulation compared to basal condition (Figure 1). Therefore, for most features, the extreme values are better biomarkers of ageing than the basal values. Virtually any evolving phenotype could be a biomarker of ageing. To determine which of these features are the best biomarkers of ageing, the Pearson’s correlation coefficients with relative age was calculated for each feature after mechanical stimulation. Out of the 1019 features used in this study, 837 have p values < 0.05 after Bonferroni correction. A list of the 50 best biomarkers and their corresponding correlation coefficients is shown in Supplementary Table 1.

**Figure 1.**
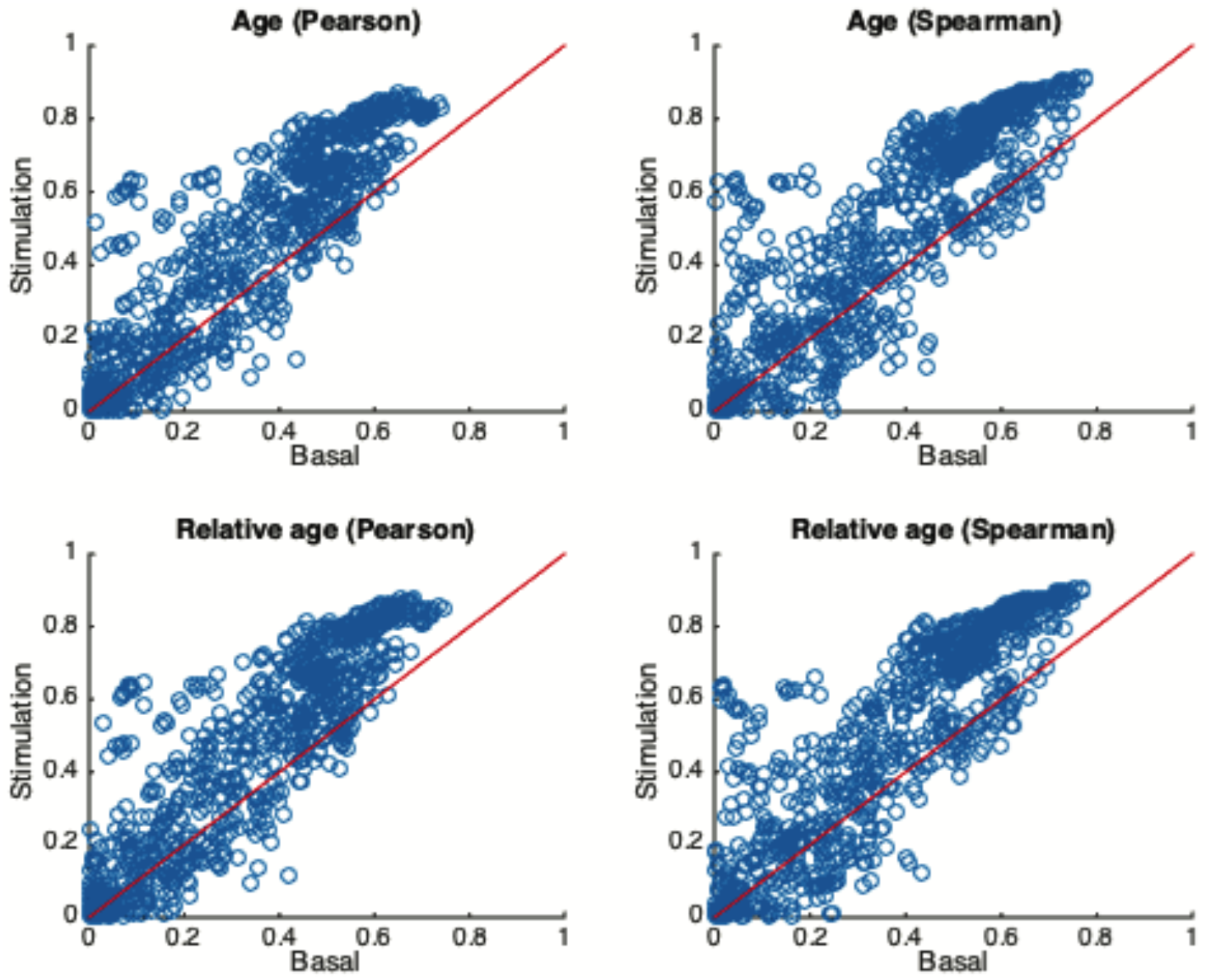
Extreme values correlate better with age and relative age. Absolute Pearson’s and Spearman’s correlation coefficients of each feature with age and relative age, obtained in basal conditions and after mechanical stimulation.

### Multidimensional phenotyping predicts age, prognosis and lifespan

Here, similarly to (Zhang et al., 2016), we use biomarkers to predict prognosis with support vector machines. In addition we predicted the age and lifespan of individual worms using a set of 100 selected features (Figure 2A). The features were selected iteratively by implementing the most predictive features one by one to determine the most predictive combination of features. As expected, the quality of the predictions increased with the number of features used before reaching a plateau. According to this result, about 100 features are sufficient to make good predictions (Figure 2B). From the best set of 100 features, age and prognosis were predicted with respective root mean standard error of 1.7 and 2.8 days while lifespan was predicted with a lower accuracy, with a root mean standard error of 3.1. This iterative approach is better predictive than the mere addition of the best predictive features (Supplementary Figure 1A). The examination of the features selected show strong divergence between the two methods (Supplementary Figure 1B). It suggests that an appropriate set of features is more informative than the mere addition of predictive features, as expected when selecting from a set of potentially correlated features.

**Figure 2.**
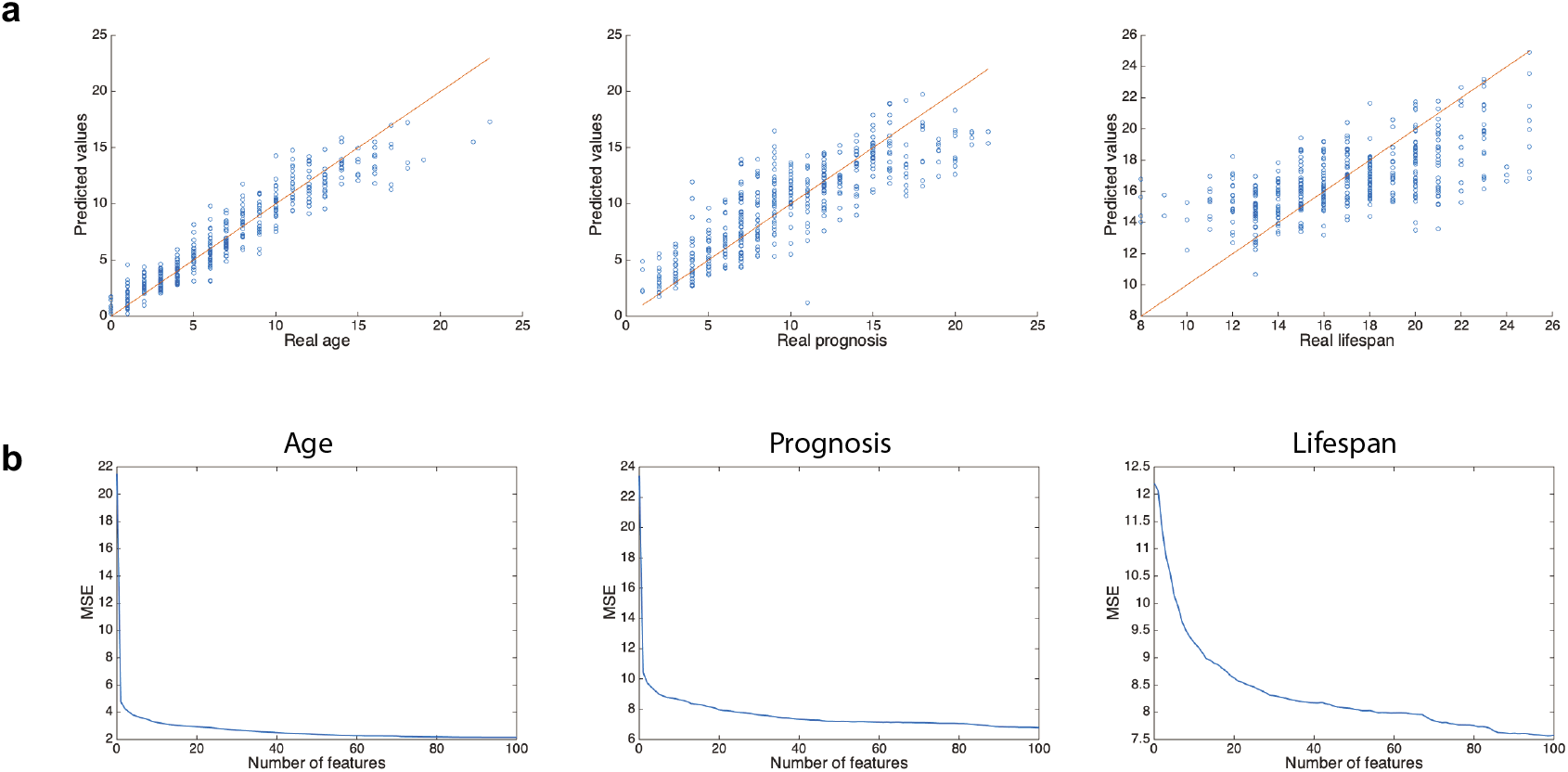
Deep phenotyping predicts age, prognosis and lifespan. (a) Predictions of age, prognosis and lifespan. Red lines indicate theoretical perfect predictions. (b) Evolution of the Mean Standard Error (MSE) over the number of features used for the predictions of age, prognosis and lifespan.

### Short-lived animals age faster than long-lived animals

Isogenic populations are composed of worms exhibiting different lifespans. This inter-individual variability between individuals in health could be explained both qualitatively, with divergent trajectories of ageing, and quantitatively, with different ageing rates. To test these two approaches, we split our cohort of 151 animals into 5 groups of different lifespans. Their phenotypes after stimulation was projected in 2 dimensions using PCA (Principal Component Analysis). The first two principal components separate individuals accordingly to age, generating a relatively well-defined trajectory of ageing. No qualitative divergence of trajectories among the lifespan groups is observed (Supplementary Figure 2). Therefore, within the same genotype, worms seem to follow a single trajectory of ageing with close starting and end phenotypes. Quantitatively, 39% of the variance of the full dataset at all ages can be described in the first principal component (Supplementary Table 2). Taking the position along the first principal component as a measure of biological age, we can use the distance between consecutive time points as an approximation of the ageing rate of the animals. From this analysis, it appears clearly that short-lived animals travel faster through the multidimensional phenotypic landscape, and therefore age faster than long-lived animals (Figure 3).

**Figure 3.**
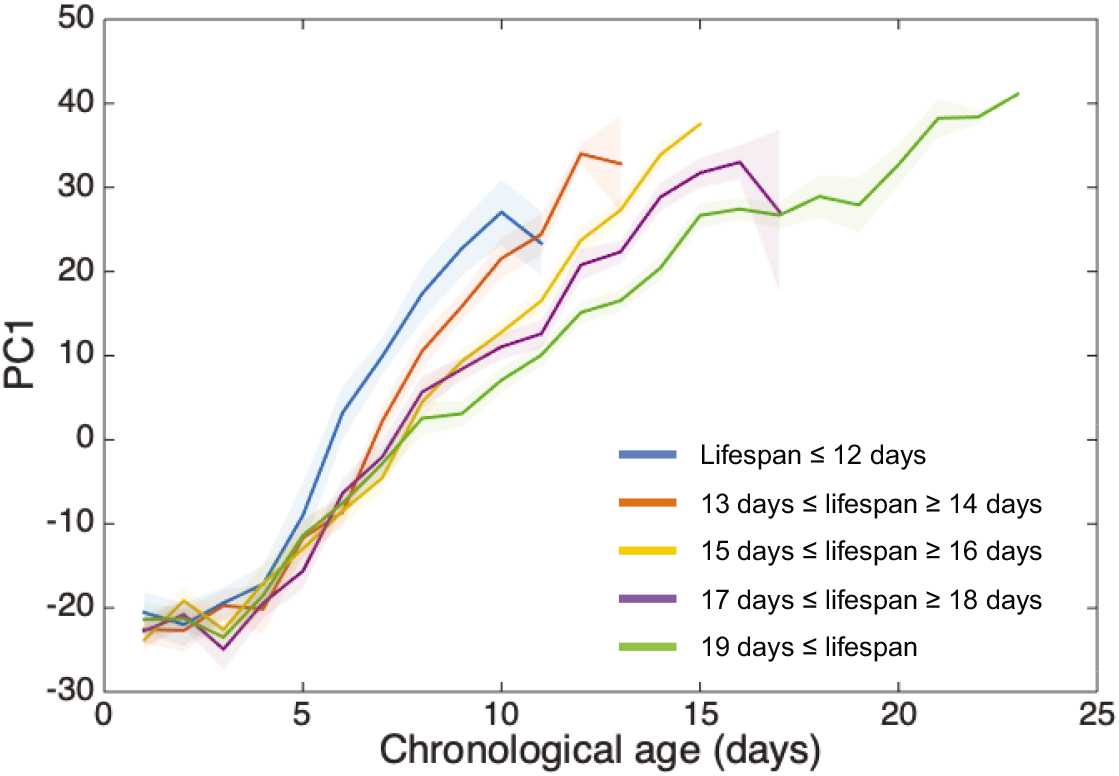
Short-lived animals age faster than long-lived animals. Evolution of the first principal component (PC1) of a principal component analysis over chronological age. Shaded areas indicate standard errors of the mean.

### Long-lived animals have a better health

Health was previously defined as a decline of maximum velocity or of prognosis (Hahm et al., 2015; Zhang et al., 2016). Here we consider health as the conservation of normal basal activity as well as the ability to reach maximum activity and defined a health index based on the range of abilities of the animal. It is calculated as the mean difference between the maximum value (90^th^ percentile) and minimum value (10^th^ percentile) for each feature and should be an indicator of the phenotypic flexibility of the animals. This index is applicable to physiological as well as pathological ageing (Martineau et al., 2019). Our health index varies along life and appeared highly dependent on lifespan. Among lifespan groups, health indexes were similar at the beginning of adulthood and decreased after day 4-5 of adulthood (Figure 4A). The rate of decline was dependent on lifespan, with a high rate of decline for short-lived animals and a low rate of decline for long-lived animals. However, in regards to relative age these rates of decline are similar, indicating that relative to lifespan, healthspans are comparable among groups (Figure 4B). Therefore, no extended twilight can be observed for long-lived worms in our conditions.

**Figure 4.**
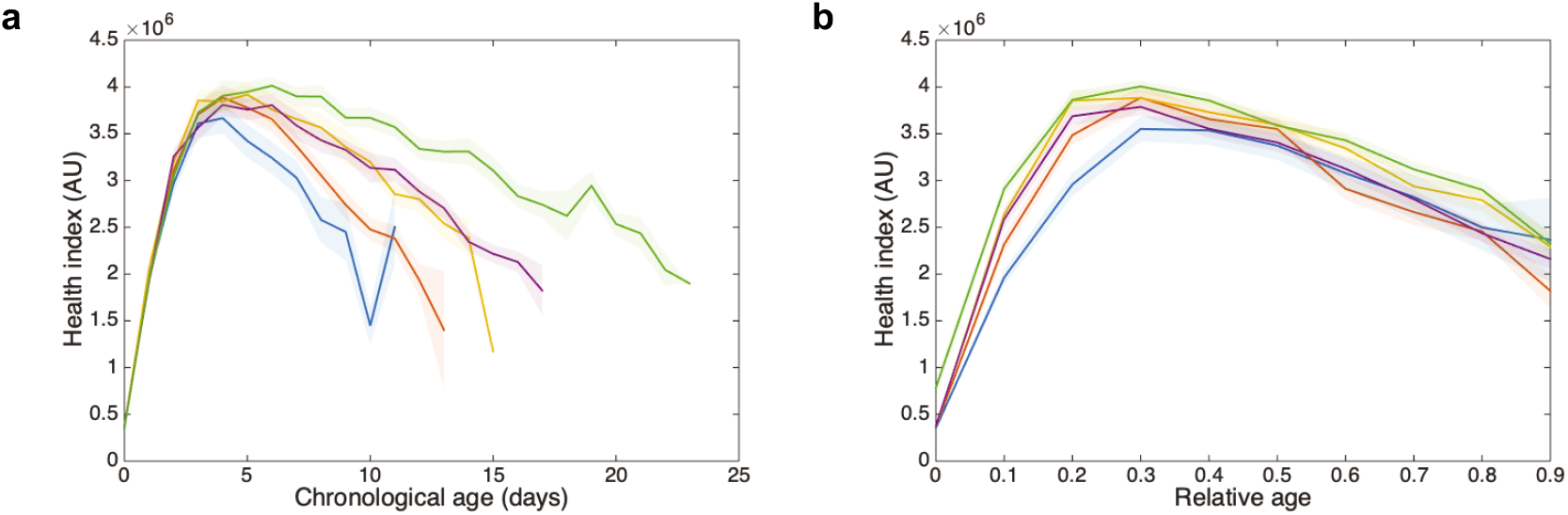
Health is coupled to lifespan. (a) Evolution of the health index over chronological age. (b) Evolution of the health index over relative age. Shaded areas indicate standard errors of the mean.

## Discussion

Prediction of age and lifespan based on health parameters is an on-going challenge of medicine. So far, human biological age can be predicted from gene expression (Peters et al., 2015), DNA methylation profiles (Hannum et al., 2013), or physical activity (Pyrkov et al., 2018), but cohorts allowing for lifespan predictions are still lacking. In *C. elegans*, individual features such as maximum velocity (at day 9 of adulthood) or a combination of a few features such as movement, autofluorescence or body size were previously shown to correlate well with lifespan or to predict prognosis (Hahm et al., 2015; Zhang et al., 2016). However, most features of the worm evolve with ageing. To better cover this wide phenotypic evolution requires a larger set of phenotypic features than previously attempted. To achieve this, we used an automated quantification of phenotypes able to describe the ageing of *C. elegans* at the organismal level with more than 1000 features. We show that this multi-parametric approach exceeds the predictive power reached with one or few pre-defined biomarkers of ageing. Indeed, the accuracy of the predictions increases with the number of features added iteratively to the model with a set of 100 non-invasive parameters appearing sufficient to achieve good predictions.

Our results show that phenotypes after mechanical stimulation better correlate with age and relative age than their counterparts obtained in basal conditions. This is consistent with previous results showing that the maximum velocity of the animals correlates better with lifespan than the mean velocity (Hahm et al., 2015). It suggests that the concept of the Short Physical Performance Battery (SPPB) can be extended to other species than human and provides support for analogies in the ageing processes between species despite extremely different morphology, postures and locomotory phenotypes (Guralnik et al., 1994).

Based on the evolution of our health index over relative age, healthspan correlates with lifespan in the wild-type strain, N2. This result differs from (Zhang et al., 2016) but confirms other previously published results (Bansal et al., 2015; Hahm et al., 2015). The type of health index, the features selected, the environment or the genotype used likely explain these discrepancies between studies. We explored many more features than previous publications (Bansal et al., 2015; Hahm et al., 2015; Zhang et al., 2016), however these features are mainly morphological and locomotory, lacking indicators for reproduction or tissue integrity, which might be crucial markers for health quantification and present in (Zhang et al., 2016). The type of set-up used for the recording of phenotypes is also important. Indeed, we recorded phenotypes in open conditions, with worms being transferred every day onto a fresh plate, avoiding the accumulation of metabolites and wastes that occur in a closed system as used in (Zhang et al., 2016). These metabolites might interfere with the good health of long-lived animals causing the observed extended twilight. Finally, the observations could be genotype-dependent, as Zhang *et al*. used a sterile strain, *spe-9(hc88)* (Zhang et al., 2016).

Health state and longevity are affected by genetics and environment. We previously observed different phenotypic trajectories of ageing for two different genotypes (Martineau et al., 2019). However even isogenic worms ageing in a controlled environment show wide inter-individual differences in lifespan. Comparing the phenotypes of short- and long-lived wild-type animals, we observe that all animals follow a similar trajectory of ageing through the multidimensional phenotypic landscape, but at different rates of ageing, similarly to (Zhang et al., 2016). Similarly to the trajectory of development, the trajectory of ageing appears robust at the organismal scale (Félix and Barkoulas, 2012; Pincus and Slack, 2010). Given the high level of stochasticity in molecular and cellular ageing observed previously (Cypser et al., 2006; Herndon et al., 2002), this observation suggests that the phenotypic repertoire of each individual evolves in a relatively coordinated mode. Nevertheless, our results show that inter-individual variability is mainly explained by the rate of ageing.

In conclusion, we demonstrated that multiple non-invasive parameters are best predictive of biological age and lifespan and are a better proxy to quantify health than single biomarker of ageing. We propose this approach as a new standard to study ageing properties at the organismal scale.

## Material and Methods

### Strains and media

The strain N2 was used in this study. Standard conditions were used to maintain and propagate this strain at 20 °C.

### Collection of behavioural data

151 single worms were video-tracked using a worm tracker equipped with a bone conductor transducer for mechanical stimulation. Worms were tracked longitudinally every day from the L4 stage to death. Worms were maintained in strict conditions at 20 °C until and during the tracking.

Single-worm tracking was performed as previously described with slight modifications. Briefly, 3 cm plates containing low peptone NGM were seeded with 20 μL of OP50 30 minutes prior tracking. Each day, each single worm was picked with a sterile eyelash on a new fleshly seeded plate and let habituate for 15 minutes. After 2 minutes of habituation in the tracker, worm behaviour was recorded for 2 minutes at 20 frames per second to extract basal phenotypes. To extract phenotypes after mechanical stimulation, behaviour was recorded for 5 seconds in basal condition before a vibration transmitted by the air of 4 seconds at 750 Hz, and recorded 1 more minute after stimulation. Videos were analysed with the freely available Tierpsy software to extract behavioural features (Javer Avelino et al., 2018). The worm-behaviour data is available on an open-source platform (Javer et al., 2018) (http://movement.openworm.org/).

### Data preparation

For each worm, a set of 4539 features were extracted with the Tierpsy software (Javer Avelino et al., 2018). These features contain information about the worm morphology, posture, locomotion and behaviour. After feature extraction, worms containing more than 25% of missing values were removed from the dataset. A selection of representative features was also operated to remove features containing missing values. The dataset was then standardised with z-score to compensate for the different units of each feature.

### Predictions

Predictions were performed with a support vector machine (linear Kernel) without cross-validation. A train set comprising 80 % of the data was used to train the model and predictions were made on the 20 % remaining data. To measure prediction accuracy over the number of features, features were tested and implemented to the model one by one, the feature giving the smallest mean standard error being selected at each iteration.

### Health index

Health index was calculated as the mean difference of maximum (90^th^ percentile) and minimum (10^th^ percentile) values for the 131 features for which percentiles were available in Tierpsy.

## Acknowledgments

CNM was the beneficiary of a fellowship from the Université Libre de Bruxelles (ULB).

## Author Contributions

Conceptualization, C.N.M., A.E.X.B., and P.L.; Methodology, C.N.M.; Formal Analysis and Investigation, C.N.M.; Resources, A.E.X.B. and P.L.; Writing – Original Draft, C.N.M.; Writing – Review and Editing, C.N.M., P.L., and A.E.X.B.; Supervision, A.E.X.B. and P.L.; Funding Acquisition, C.N.M.

