## Supplementary material for "Multidimensional phenotyping predicts lifespan and quantifies health in *C. elegans*": Tables S1 and S2, Figures S1 and S2

| Features | Correlation coefficient |
|---|---|
| 'd_curvature_mean_hips_abs_90th' | 0,879 |
| 'angular_velocity_hips_abs_90th' | 0,875 |
| 'relative_to_head_base_angular_velocity_head_tip_abs_90th' | 0,871 |
| 'area_90th' | 0,869 |
| 'angular_velocity_tail_base_abs_90th' | 0,867 |
| 'curvature_tail_IQR' | 0,863 |
| 'curvature_std_neck_50th' | 0,863 |
| 'd_relative_to_body_angular_velocity_tail_tip_50th' | 0,863 |
| 'd_relative_to_hips_radial_velocity_tail_tip_90th' | 0,862 |
| 'curvature_std_neck_IQR' | 0,861 |
| 'curvature_tail_50th' | 0,860 |
| 'relative_to_tail_base_angular_velocity_tail_tip_abs_IQR' | 0,860 |
| 'd_relative_to_body_angular_velocity_tail_tip_IQR' | 0,860 |
| 'angular_velocity_abs_IQR' | 0,859 |
| 'd_curvature_std_tail_abs_IQR' | 0,859 |
| 'curvature_neck_IQR' | 0,858 |
| 'curvature_neck_50th' | 0,856 |
| 'angular_velocity_tail_base_abs_IQR' | 0,855 |
| 'd_width_head_base_90th' | 0,855 |
| 'd_speed_hips_90th' | 0,854 |
| 'minor_axis_IQR' | 0,854 |
| 'minor_axis_90th' | 0,852 |
| 'eigen_projection_7_IQR' | 0,851 |
| 'd_relative_to_tail_base_radial_velocity_tail_tip_90th' | 0,851 |
| 'curvature_mean_neck_IQR' | 0,851 |
| 'angular_velocity_head_tip_abs_IQR' | 0,851 |
| 'eigen_projection_6_50th' | 0,850 |
| 'eigen_projection_6_IQR' | 0,850 |
| 'curvature_mean_neck_50th' | 0,850 |
| 'eigen_projection_7_50th' | 0,850 |
| 'relative_to_body_speed_midbody_abs_IQR' | 0,849 |
| 'speed_IQR' | 0,844 |
| 'path_density_tail_95th' | 0,844 |
| 'eigen_projection_5_IQR' | 0,844 |
| 'curvature_std_head_50th' | 0,843 |
| 'd_curvature_tail_IQR' | 0,843 |
| 'd_curvature_tail_50th' | 0,843 |
| 'd_curvature_std_neck_50th' | 0,842 |
| 'path_transit_time_tail_50th' | 0,842 |
| 'd_curvature_mean_midbody_abs_90th' | 0,841 |
| 'd_curvature_neck_50th' | 0,841 |
| 'eigen_projection_5_50th' | 0,841 |
| 'eigen_projection_2_IQR' | 0,841 |
| 'd_speed_midbody_IQR' | 0,841 |
| 'angular_velocity_tail_tip_abs_90th' | 0,841 |
| 'd_path_curvature_body_abs_90th' | 0,840 |
| 'eigen_projection_6_abs_IQR' | 0,840 |
| 'd_length_90th' | 0,840 |
| 'd_curvature_neck_IQR' | 0,840 |
| 'curvature_std_tail_50th' | 0,838 |

**Table S1. Biomarkers of ageing.**

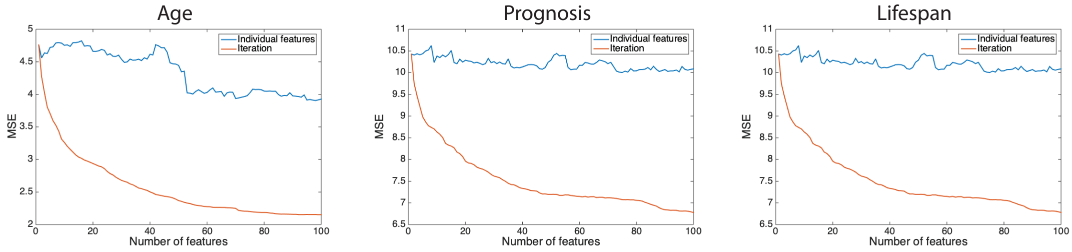


**Figure S1. Deep phenotyping predicts age, prognosis and lifespan.** (a) Evolution of the Mean Standard Error (MSE) over the number of features used for the predictions of age, prognosis and lifespan.

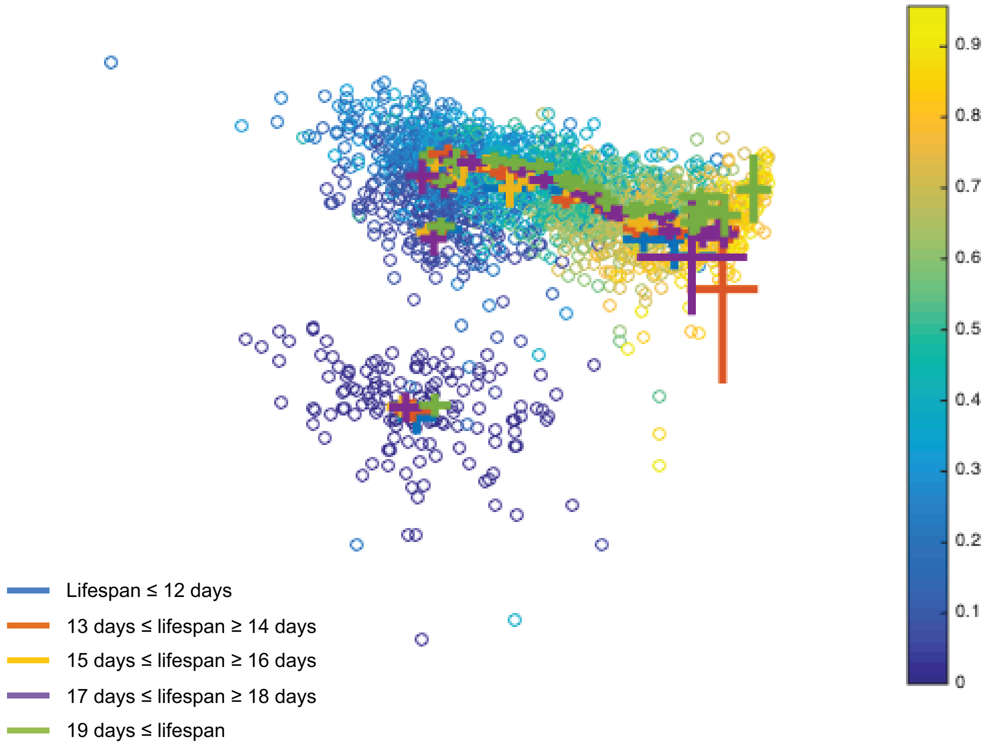


**Figure S2. No phenotypic divergence among lifespan groups.** PCA representation of the phenotypes of 5 different lifespan groups over age. Color of each dot indicates the relative age, from L4 (dark blue) to death (yellow). Crosses indicate mean phenotype and standard error of the mean for each age.

| Principal components | % Variance explained |
|---|---|
| 1 | 39,18 |
| 2 | 10,14 |
| 3 | 4,10 |
| 4 | 3,79 |
| 5 | 2,82 |
| 6 | 2,14 |
| 7 | 1,83 |
| 8 | 1,53 |
| 9 | 1,38 |
| 10 | 1,27 |

**Table S2. Percentage of variance explained by the 10 first principal components.**
